# Mechanistic Insights into TYSM Protein Regulation by mTORC2 in Response to Chemotherapy

**DOI:** 10.1101/2025.05.06.652447

**Authors:** Abdulrahman El Sayed, Maciej Zakrzewski, Remigiusz Serwa, Abdelhalim Azzi

## Abstract

Transcriptional and translation control of thymidylate synthase (TYMS) is poorly understood, particularly in response to chemotherapeutic drugs such as 5-Fluorouracil (5-FU) and its derivatives. The current study addressed this gap by demonstrating a biphasic response in TYMS protein levels upon 5-FU treatment. Indeed, we observe an initial reduction within the first few hours, followed by a marked increase at 24 hours. These changes occurred independently of transcriptional regulation, as TYMS mRNA levels remained stable during the early phase and showed only a moderate increase later. We further showed that neither autophagy nor proteasomal degradation contributed to this dynamic, but instead it is driven by change in its translation. Using thermal proteome profiling, we identified SIN1, a key component of the mTORC2 complex, as a key regulator of TYMS protein levels. Functional studies revealed that SIN1 depletion negatively alters TYMS levels and dynamics and sensitizes cancer cells to 5-FU-mediated cell death. These findings uncover a novel mechanism controlling TYMS protein levels and suggest that targeting SIN1 may represent a promising strategy to enhance the therapeutic efficacy of 5-FU-based treatments.

## Introduction

Thymidylate synthase (TYMS) plays a key role in DNA synthesis. TYMS protein catalyzes the conversion of deoxyuridine monophosphate (dUMP) to deoxythymidine monophosphate (dTMP), playing a central role in maintaining nucleotide pools necessary for cell proliferation [1-2]. TYMS RNA and protein levels were found to be significantly up-regulated in various cancers and associated with the poor prognosis of patients [3-4]. Given its essential role in DNA replication, TYMS is considered as a primary target of chemotherapeutic agents developed to inhibit nucleotide biosynthesis in rapidly proliferating cancer cells [5-6]. One of the most widely used inhibitors of TYMS is 5-fluorouracil (5-FU). Upon entry into the cell, 5-FU is converted into metabolites: fluorodeoxyuridine monophosphate (FdUMP), fluorodeoxyuridine triphosphate (FdUTP), and fluorouridine triphosphate (FUTP). Incorporation of these metabolites during DNA and RNA synthesis leads to DNA and RNA damage and death of rapidly proliferating cells. Moreover, the binding of 5-FdUMP to TYMS leads to the formation of an inactive complex, altering TYMS activity, which in turn causes deoxynucleotide (dNTP) imbalance and DNA damage [7]. Despite the widespread clinical use of 5-FU and its derivatives, resistance to treatment remains a significant challenge. Indeed, studies have shown that prolonged treatment with 5-FU or its derivative leads to a marked increase in TYMS protein levels [8-9]. The exact mechanism behind this increase is still not clear. TYSM is an RNA-binding protein that interacts with its own mRNA as well as others, such as P53, thereby regulating both its own and other protein translation. Upon interaction with FdUMP, the RNA binding capacity of TYMS is reduced, thus enhancing its own translation [10-12]. Whether this is the only mechanism that controls TYMS protein dynamics is currently unknown. Our study addresses this gap by examining how 5-FU and its derivatives influence TYMS protein dynamics. We have shown that TYMS protein exhibits a biphasic response, with TYMS protein levels significantly reduced within the first few hours of treatment, followed by a marked increase at 24 hours. These changes occurred independently of transcriptional changes, as TYMS mRNA levels remained stable in the early phase of treatment and showed only a moderate increase at later time points. Moreover, neither autophagy nor proteasomal degradation contributed to this dynamic. Using thermal proteome profiling, we identified MAPKAP1 (SIN1 protein), a key component of the mTORC2 complex, as a key regulator of TYMS levels. Using a small hairpin targeting the SIN1 protein, we showed that TYMS protein dynamics following 5-FU or FuDR treatment required intact SIN1 protein. Furthermore, SIN1 depletion sensitizes cancer cells to 5-FU and FuDR-mediated cell death. These findings highlight a novel mechanism of TYMS regulation and suggest that targeting SIN1 may provide new therapeutic avenues to modulate TYMS levels and enhance the efficacy of 5-FU treatments.

## Results

### TYMS Protein levels exhibit biphasic response to 5-FU and its derivatives

To examine the immediate cellular response to 5-fluorouracil (5-FU), we used two breast cancer cell lines, MCF7 and MDA-MB-231. We treated these cell lines with 100 µM 5-FU for a short duration (2, 4, and 6 hours). Interestingly, our data revealed that short treatment of cells with 5-FU led to a significant decrease in thymidylate synthase (TYMS) protein levels in both cell lines (**Figure 1 A-D**). These changes also occur at lower concentrations of 5-Fu (**Sup. Figure 1 A-D**). We next examined the impact of longer treatment with 5-FU. Our data revealed that prolonged treatment of cells with 5-FU led to a significant increase in the levels of thymidylate synthase (TYMS) protein in both cell lines, suggesting that 5-FU metabolites have a similar effect on cells irrespective of the origin of breast cancer cells (**Figure 1 E-G**). Moreover, our data are in line with previous findings showing an increase in TYSM levels following prolonged treatment with 5-FU in colorectal cancer cells [8, 14].

**Figure 1.**
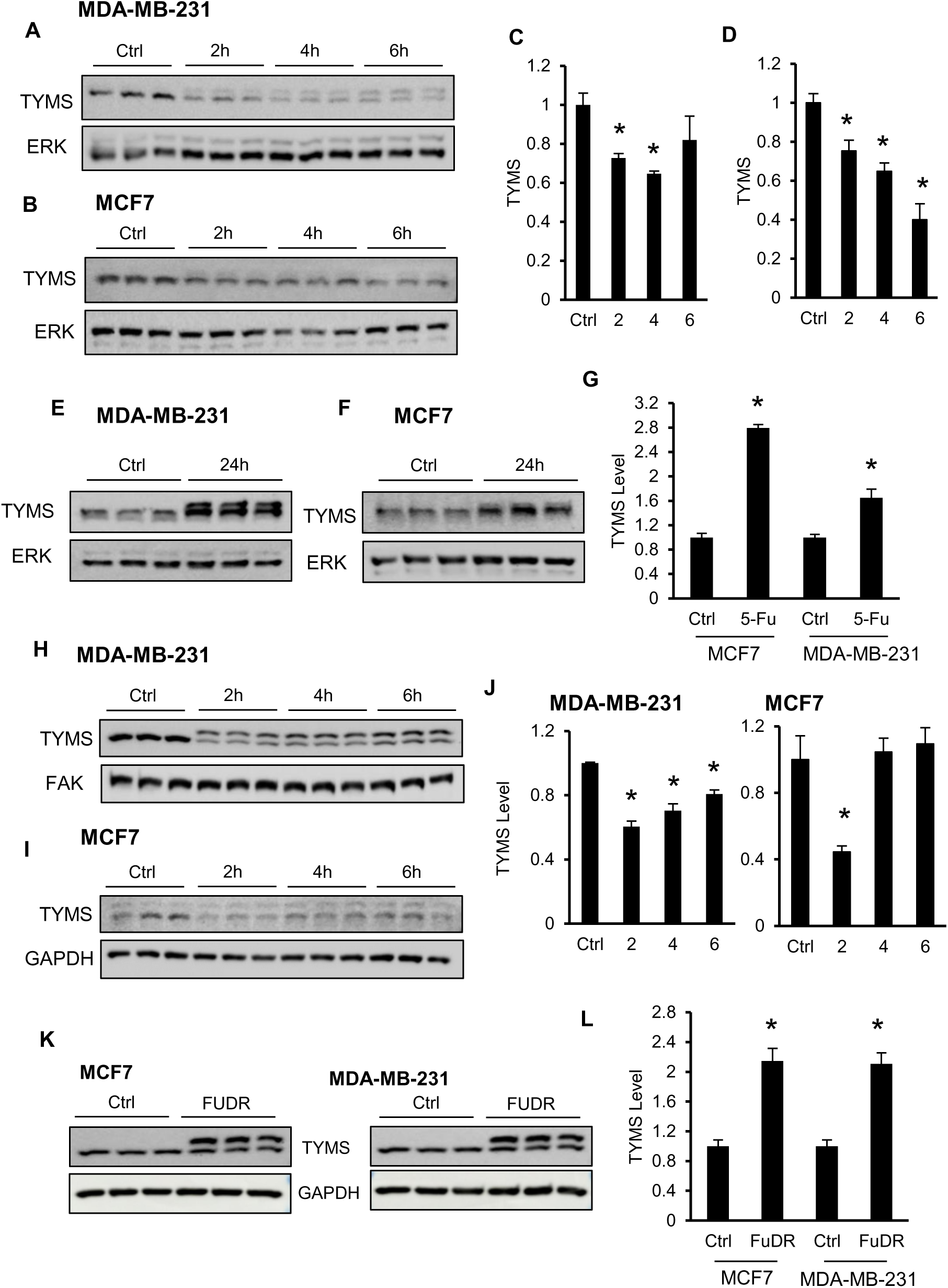
5-fluorouracil and derivatives alter TYMS protein level time-dependent manner. **(A-B)** Lysates from MDA-MB-231 and MCF-7 cells treated with 100 µM 5-FU for the indicated durations (h: hours) and then stained with the indicated antibodies. **(C-D)** Bar graphs showing the results of densitometric analysis of TYMS protein levels under different conditions obtained in A and B, respectively. The data are presented as the means ±SEMs. ANOVA, multiple comparisons: Dunnett test. MDA-MB-231: F (3, 8) = 4.809, P <0.05. MCF7: (F (3, 8) = 17.83, P <0.001). **(E, F)** Lysates from MDA-MB-231 and MCF-7 cells treated with 100 µM 5-FU for 24 hours and stained with the indicated antibodies. **(G)** Bar graphs showing the results of densitometric analysis of TYMS protein levels under different conditions obtained in E and F. Unpaired T test, **, P˂0.01. **(H-I)** Lysates from MDA-MB-231 and MCF-7 cells treated with 50 µM FuDR for the indicated durations and then stained with the indicated antibodies. **(J)** Bar graphs showing the results of densitometric analysis of TYMS protein levels under different conditions obtained for each cell line in H and I, respectively. The data are presented as the means ±SEMs. ANOVA, multiple comparisons: Dunnett test. MDA-MB-231: F (3, 8) = 30.35, P<0.001. MCF7: F (3, 8) = 9.587, P<0.001). (**K-L**) Lysates from MDA-MB-231 and MCF-7 cells treated with 50µM FuDR for 24 hours and then stained with the indicated antibodies. The bar graphs show the results of densitometric analysis of TYMS protein levels under different conditions obtained for each cell line. The data are presented as the means ±SEMs. Unpaired T test, **, P˂0.001.

5-FU is converted into the nucleoside metabolites fluorodeoxyuridine monophosphate (FdUMP) and fluorodeoxyuridine triphosphate (FdUTP). Subsequent metabolism of FdUMP and FdUTP leads to DNA and RNA damage, respectively. FdUMP can also bind and inhibit thymidylate synthase (TYMS) [2,7]. Next, we used 5-fluorouridine (FUR) and 5-fluorodeoxyuridine (FuDR), which are intracellularly metabolized and converted into FdUTP and FdUMP, respectively. Surprisingly, short-duration treatment of cells with FuDR and FuR similarly alters TYMS protein, with a decrease at early and a significant increase at later time points (**Figure 1 H-L and Sup. Figure 2 A-D**). To rule out the possibility that the decrease in TYMS levels at the early time point is a result of a change in its thermodynamics upon binding to 5-FU or its derivatives, a drug affinity responsive target stability assay (DART) was carried out to examine the binding of TYMS to FUR and FuRD [15]. This assay confirmed the direct interaction between TMYS and FuDR, significantly decreasing TYMS protein. In contrast, FuR had no effect, demonstrating that a decrease in TYMS protein upon treatment with 5-FU or derivatives does not result from drug-protein interaction (**Sup. Figure 2 E-F)**. Together, these data highlight time-dependent changes in TYMS protein upon treatment with pyrimidine analogues.

### Autophagy and proteasome inhibition do not alter TYMS protein levels following 5-FU Treatment

The dynamic changes in TYMS protein level upon 5-FU treatment might be driven by (**I**) changes in protein stability, (**II**) changes in protein translation, or (**III**) alterations in the transcription level of the gene encoding TYMS protein. We thus analyzed the mechanism that drives the decrease in TYMS protein upon short treatment with 5-FU. We used proteasome inhibitor bortezomib or Mg132 and the autophagy inhibitors bafilomycin or chloroquine to examine changes in TYMS protein degradation [16-17]. Simultaneous treatment of cells with 5-FU and these inhibitors did not interfere with the decrease in TYMS protein levels but promoted a further decrease of the protein, especially in cells treated with chloroquine (**Figure 2 A-B and Sup. Figure 3 A-B**). These results suggest that alterations in TYMS protein could result from changes in the transcriptional activity of the gene encoding the protein. To test this possibility, we analyzed the RNA encoding TYMS upon short and prolonged treatment with 5-FU or FuDR. Interestingly, this analysis showed no changes in TYMS RNA levels after 2 hours of treatment with 5-FU or FuDR, and only a moderate increase occurred after 24 hours, suggesting that the decrease seen in TYMS protein levels at early hours could be a result of changes in translation (**Figure 2 C-D)**. To examine this possibility, we used the threonine analogue, β-ethynylserine (βES) for metabolic labelling of the newly synthetized proteins. When combined with click chemistry and enrichment methods, this approach enable specific detection of nascent proteins within a minutes. As indicated in Figure 2E, in vehicle-treated cells, newly synthesized TYMS protein was detected within two hours. In contrast, TYMS was nearly undetectable in cells treated with FuDR or Fur, demonstrating that the decrease in TYMS protein within the first two hours of treatment with drugs is a result of the change in its translation.

**Figure 2.**
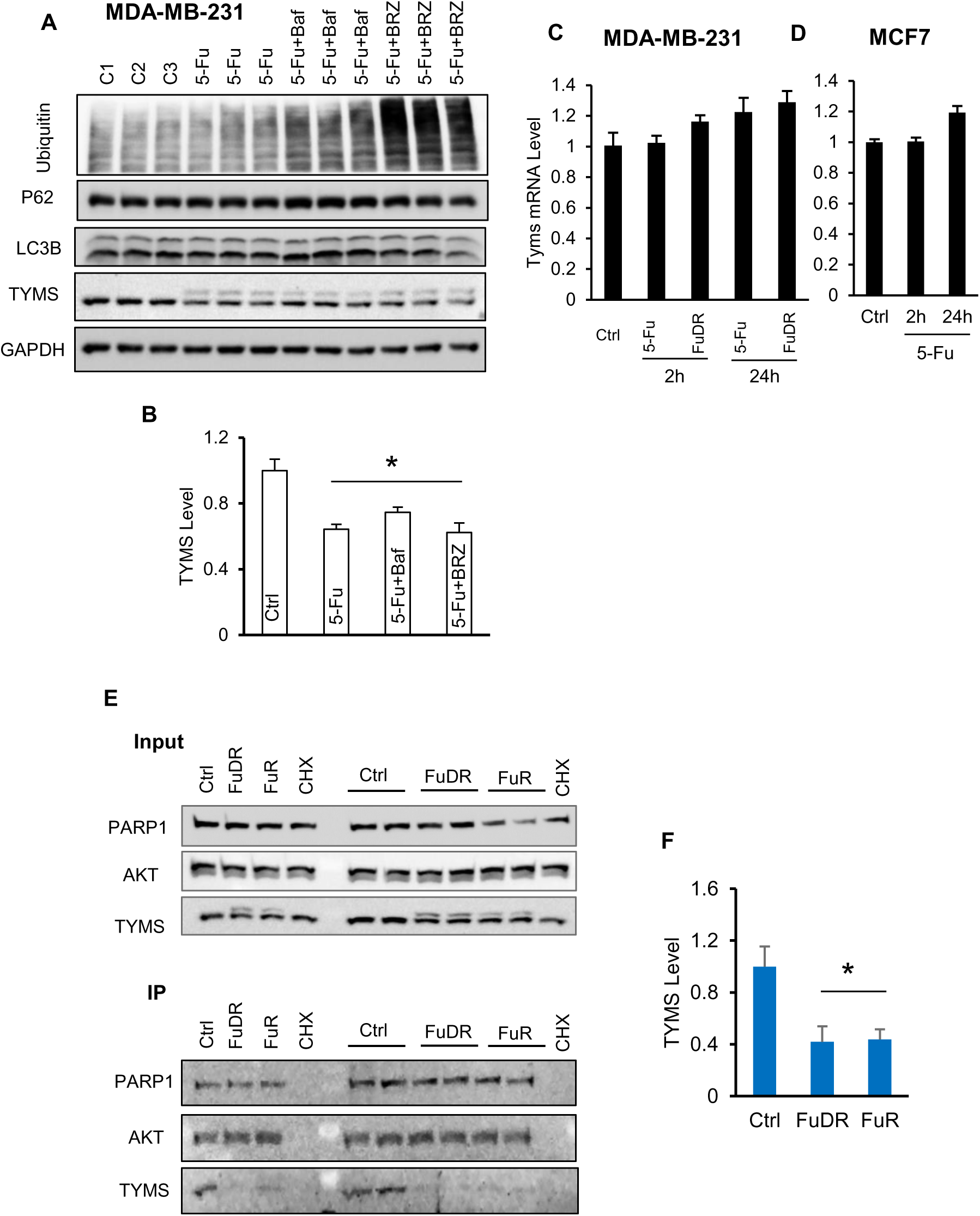
5-fluorouracil-mediated TYMS decrease is driven by a change in translation. **(A)** Lysates from MDA-MB-231 cells pre-treated with 10 nM bafilomycin A or 100 nM bortezomib for 2 hours, followed by 100µM 5-FU for 2 hours. **(B)** Bar graphs showing TYMS protein levels under different conditions obtained in A. The data are presented as the means ±SEMs. ANOVA, multiple comparisons: Dunnett test. F (3, 8) = 11.97, P˂0.005. **(C-D)** Bar graphs showing Tyms mRNA levels following treatment with 100 µM 5-FU for the indicated durations. The data are presented as the means ±SEMs. ANOVA, multiple comparisons: Dunnett test. **MCF7:** F (2, 8) = 21.91, P˂0.005. MDA-MB-231: 5-FU (F (2, 8) = 2.48, P=0.163), FuDR (F (2, 8) = 4.25, P=0.07). (**E**) Visualization of newly synthesized protein in MDA-MB-231 cells treated or not with 50 µM FuDR or 50 µM FuR during two hours, combined with β-ethynylserine (βES). Gels show the total input (5%) and the newly synthesized proteins after enrichment. PARP1 and AKT were used as internal controls. (**F**) The bar graph shows the amount of newly synthesized TYSM protein under the conditions shown in E.

### The PI3K/AKT/mTOR signalling pathway exhibits a biphasic response to 5-FU and its derivatives

The mTOR signalling pathway is a central regulator of cell growth and metabolism. This pathway exists under two protein complexes, mTORC1 and mTORC2. mTORC1 is an energy sensor and controls protein translation, whereas mTORC2 controls cell proliferation and cytoskeletal remodelling [18]. Studies have shown that inhibition of the PI3K/Akt/mTOR signaling pathway enhances the antitumor effect of 5-FU [19-20]. However, the dynamic activity and the contribution of this signaling pathway to 5-FU resistance remain poorly described.

Using phosphorylated P70S6 kinase at Thr389 and phosphorylated AKT1 at Ser473 as a readout of mTORC1 and mTORC2 activity [21], respectively, we analyzed the impact of the 5-Fu or its derivative on the phosphorylation levels of these two proteins. In both cell lines, 5-FU treatment led to a marked decrease in phospho-p70 S6 kinase levels, with a marked effect in MCF7 cells. In contrast, the same treatment moderately increased AKT1 Ser473 phosphorylation in both cell lines (**Sup. Figure 4 A-D**). Consistently, short-duration treatment of cells with FuDR leads to similar changes in mTORC1 and mTORC2 activity, albeit with different magnitudes in the analyzed cell lines, highlighting a cell type-dependent effect of 5-FU and its derivative on the mTORC activity (**Sup. Figure 4 E-H**).

Given the role of mTORC in translation [21], these findings suggest that the decrease in TYMS level upon short treatment with 5-FU and derivative may be directly linked to a reduction of mTORC1 activity. To test this hypothesis, we first examined the impact of a panel of inhibitors targeting PI3K/AKT or mTORC1. As indicated in supplementary figure 5, application of PI3K/AKT inhibitor did not affect the basal TYMS protein levels. In contrast, prolonged treatment of cells with the mTORC inhibitor rapamycin led to a marked decrease in TYMS protein. Short-term treatment of cells with rapamycin primarily affects mTORC1, whereas prolonged treatment inhibits mTORC1 and mTORC2 [22]. Therefore, our data suggest that the observed change in TYMS dynamics may be mediated by mTORC2.

Next, we analyzed the role and activity of mTORC1/2 upon prolonged treatment with FuDR. Consistent with the above results, P70S6 kinase phosphorylation is significantly lower following prolonged treatment with FuDR, whereas AKT1 at Ser473 remains unchanged, highlighting dynamic changes in mTORC2 activity over time (Figure **3 A, D**). Moreover, this experiment confirmed that rapamycin alone significantly alters TYMS protein levels. When combined with FuDR, a marked decrease in TYMS levels is also observed in MCF7 cells. Nevertheless, TYMS protein levels remain higher in MDA-MB-231 cells simultaneously treated with rapamycin, suggesting that these cells may resist lower rapamycin concentrations upon stress (Figure **3A, D**). Furthermore, a further increase in AKT1 phosphorylation following prolonged treatment with rapamycin was not accompanied by a marked increase in TYMS, suggesting that changes in TYMS protein are independent of the PI3/AKT signalling axis. Moreover, these data highlight an additional separate mechanism that controls TYMS protein.

**Figure 3.**
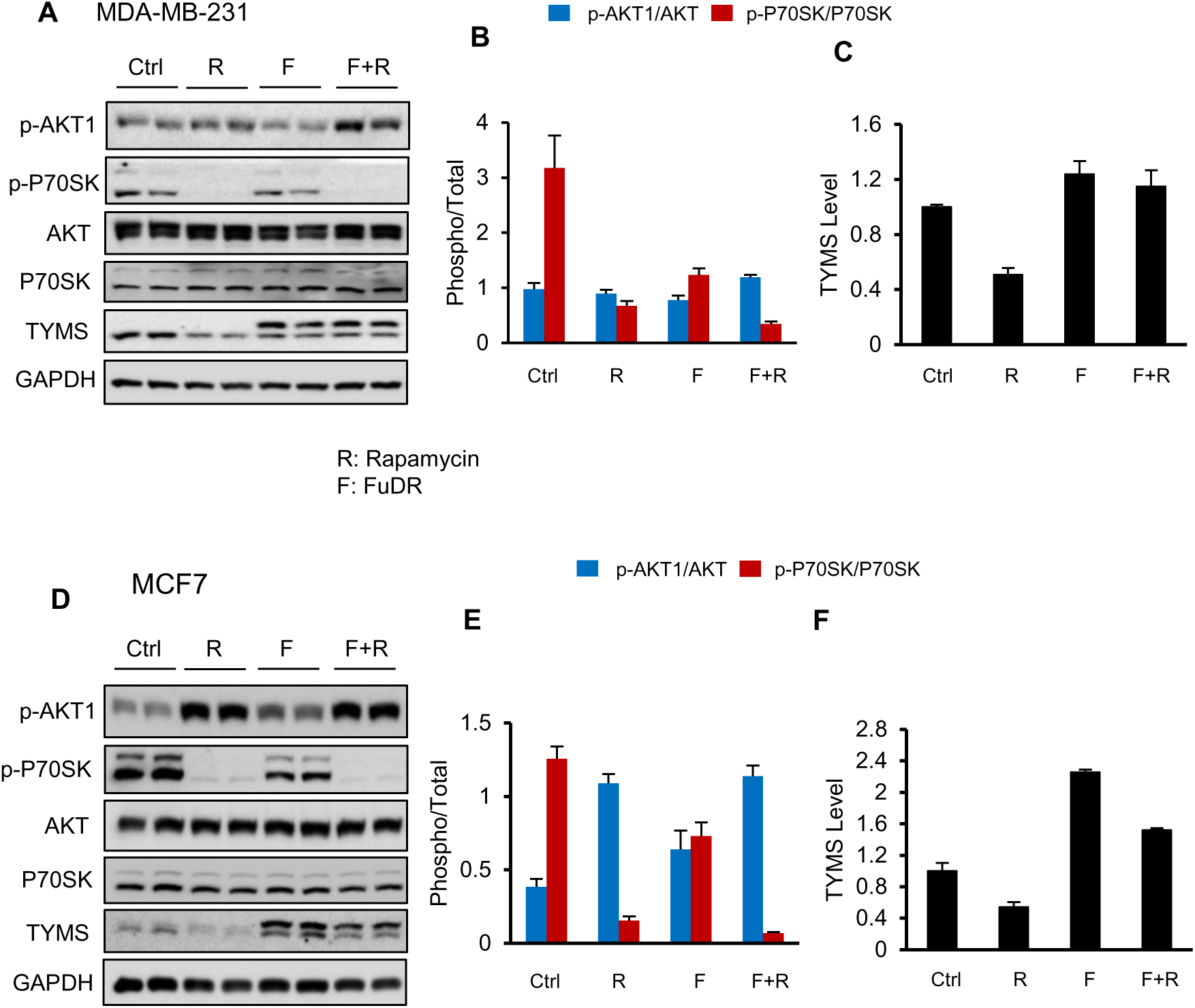
Changes in TYMS levels are mTORC1 independent. **(A)** Lysates from MDA-MB-231 cells treated or not with 50 µM 5-FuDR or 50 nM Rapamycin for 24 hours were stained with the indicated antibodies. **(B-C)** Bar graphs showing the results of densitometric analysis of the protein levels under different conditions obtained in A. The data are presented as the means ±SEMs. ANOVA, multiple comparisons: Dunnett test. p-AKT1 (F (3, 8) = 4.06, P<0.05), p-P70SK (F (3, 8) = 17.5, P<0.001), TYMS (F (3, 8) = 16.34, P<0.001). **(D)** Lysates from MCF7 cells treated or not with 50µM 5-FuDR, 50nM Rapamycin for 24 hours were stained with the indicated antibodies. **(E-F)** Bar graphs showing the results of densitometric analysis of the protein levels under different conditions obtained in D. The data are presented as the means ±SEMs. ANOVA, multiple comparisons: Dunnett test. p-AKT1 (F (3, 8) = 18.5, P<0.0005), p-P70SK (F (3, 8) = 76.9, P<0.0001), TYMS (F (3, 8) = 124.1, P<0.0001).

**Figure 4.**
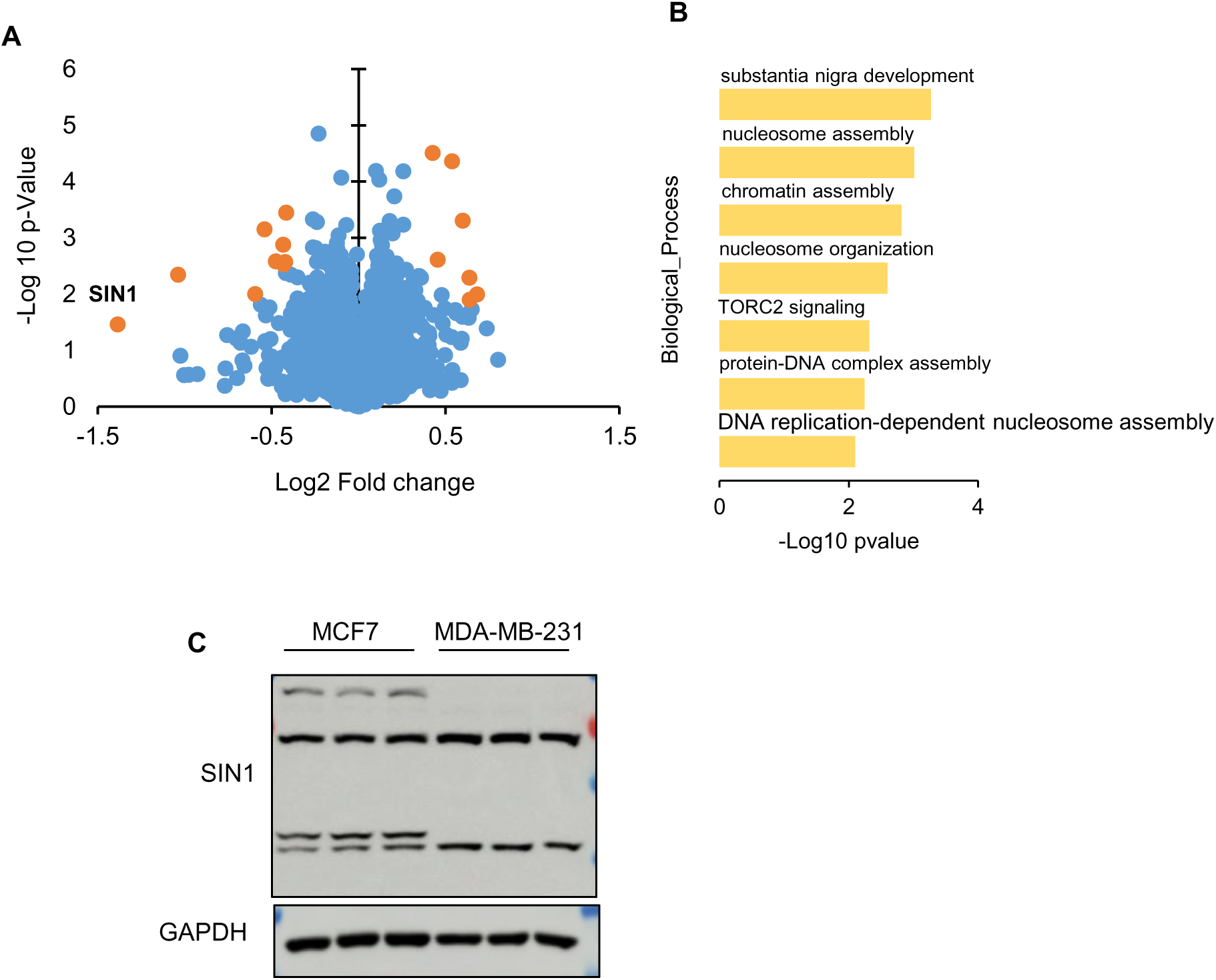
Proteome Integral Solubility Alteration analysis of 5-fluorouracil in MDA-MB-231 cells. (**A**) Volcano plot of PISA results expressed as log-transformed values of the ratio of protein soluble amount per protein of 5-FU-treated compared to control-treated cells, and relative p-value. Each dot represents individual proteins identified in the final analysis. The most significant hits are labeled with in orange font (FDR ≤ 0.05). Bleu dots represent identified proteins with no significant change (FDR p > 0.05). (**B**) Bar graph showing log10 p-value and GO terms of the biological processes affected by the significant identified hits using EnrichR KG. (**C**) Western blot analysis of SIN1 isoform level in MCF7 and MDA-MB-231. 3 biological replicates.

### Thermal proteome profiling reveals a potential role for mTROC2 in cellular response to 5-FU

To screen for potential targets of 5-fluorouracil (5-FU), we used the Proteome Integral Solubility Alteration (PISA) assay. This experimental approach allows the identification of protein-drug interactions based on thermal stability changes in the proteome upon drug binding, preserving native cellular contexts [23]. Given the similar changes in TYMS levels with both FuR and FuRD, we decided to use 5-FU to examine its early effect on the proteome of MDA-MB-231 cells. Using PISA methods, cells were first treated with 200 µM 5-FU or vehicle for 2 hours to capture early cellular changes of the proteome under very low intracellular concentrations of FuDR and FuR, assuming a slow conversion for 5-FU into FuDR and FuR [24]. Consistently, quantitative mass spectrometry analysis showed minimal changes in the thermal stability of the proteome. Indeed, compared to vehicle-treated cells, only 16 proteins showed significant changes in thermal stability, of which 9 showed lower, and 7 showed higher thermal stability upon 5-FU treatment (FDR p-value <0.05) (**Figure 3 A**). Despite the low number of significant candidates, pathway analysis showed that thermally affected proteins are mainly involved in protein-DNA interactions (**Figure 3 B**), which is in line with the recent observation reported with FuDR [25]. Interestingly, the mTORC2 signaling pathway is also affected. Indeed, the SIN1 protein (MAPKAP1), a component of the mTROC2 complex, displays a significant decrease in thermal stability (**Figure 3 A**). As an integral part of the TORC2 complex, SIN1 protein has been described to regulate several processes, including response to growth factors, response to genotoxic stress, and DNA damage. Moreover, studies have shown that this protein has several transcript isoforms encoding various proteoforms [13]. Consistently, our data show that MCF7 and MDA-MB-231 express different levels of each SIN1 isoform (**Figure 3 C**). The exact function of each isoform in regulation of mTORC2 activity and cellular response to stress stimuli is not well known. For instance, preliminary studies have shown that some isoforms play a key role in mTOCR2 signaling, whereas others could regulate separate biological processes [13, 26].

### SIN1 protein is required for cellular response to 5-FU metabolite and protects cells from genotoxic stress

Our data above showed that SIN1 is among the top proteins whose function could be immediately affected at the early point of treatment with 5-Fu or its derivatives. Previous studies have shown that SIN1 is required for mTORC2 complex assembly and AKT-Ser473 phosphorylation in response to growth factors and DNA-damaging agents [27-28]. To examine the role of SIN1 in response to pyrimidine analogues, we used a lentiviral knockdown method to achieve a stable reduction in the RNA encoding SIN1 in MCF7 and MDA-MB-231. As indicated in the supplementary figure 6, this approach significantly reduces RNA encoding SIN1 and all its proteoforms. Analysis of mTORC1/2 activity showed that basal mTORC1 is slightly higher in Sh-SIN1 cells, as seen with the increase in P70S6 kinase phosphorylation. In contrast, the inverse occurs for mTORC2, as seen with markedly lower levels of AKT1 Ser473 phosphorylation. Interestingly, our data show that SIN1 depletion leads to low levels of TYMS protein, and short-term treatment of cells with FuRD led to almost undetectable levels of this protein in these cells, highlighting a role for SIN1 in the control of TYMS (**Sup. Figure 6 D-E**).

Long-term treatment of cells with FuRD led to the complete inability of Sh-SIN1 cells to activate mTORC2, as seen with markedly low levels of AKT1 phosphorylation. Moreover, this analysis also showed that long term stress induced increase in the level of TYMS protein are markedly altered in cells in which SIN1 was knocked down, suggesting a key role for SIN1 in regulating TYMS protein under both basal and stress conditions. Furthermore, SIN1 seems to be required for WT P53 stability and activity, as shown by lower P53 and P21 protein levels in Sh-SIN1 treated cells with FuDR (**Figure 5A-I**).

**Figure 5.**
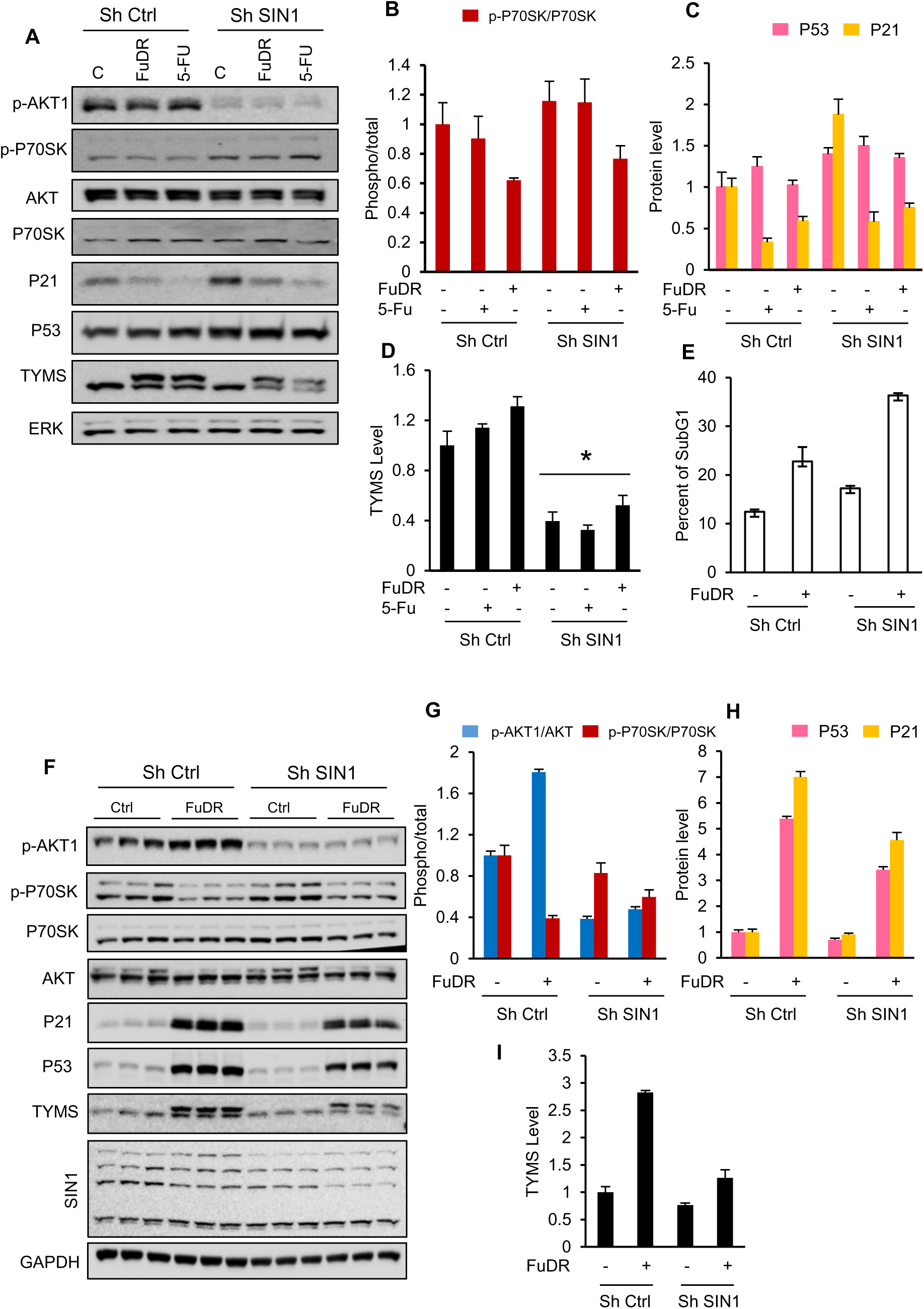
SIN1 controls AKT activity and TYMS levels following genotoxic stress. **(A)** Lysates from MDA-MB-231 cells stably expressing Sh control or small hairpin against SIN1, treated or not with 50µM 5-FuDR or 100µM 5-FU during 24 hours, were stained with the indicated antibodies. **(B-D)** Bar graphs showing the results of densitometric analysis of the protein levels under different conditions obtained in A. The data are presented as the means ±SEMs. ANOVA, multiple comparisons: Dunnett test. p-P70SK (F (5, 12) = 2.864, P=0.0628), P53 (F (5, 12) = 3.498, P<0.05, P21 (F (5, 12) = 26.09, P<0.0001). TYMS (F (5, 12) = 32.68, P<0.0001). **(E)** Bar graphs showing the percentage of Sh Ctrl and Sh SIN1 cells at SubG1 after 72 hours of treatments with FuDR. Data are presented as the means ±SEMs. ANOVA, (F (3, 8) = 45.1, P<0.0001). **(F)** Lysates of MCF7 cells stably expressing Sh control or small hairpin against SIN1 treated or not with 50µM 5-FuDR during 24 hours stained with the indicated antibodies. **(G-I)** Bar graphs showing the results of densitometric analysis of the protein levels under different conditions obtained in F. The data are presented as the means ±SEMs. ANOVA, multiple comparisons: Dunnett test. p-AKT1 (F (3, 8) = 483.1, P<0.0001), p-P70SK (F (3, 8) = 11.58, P<0.01), P53 (F (3, 8) = 616.3, P<0.0001), P21 (F (3, 8) = 251.3, P<0.0001), TYMS (F (3, 8) = 97.82, P<0.0001).

Having shown the crucial role of SIN1 in mTORC2/AKT activation and control of TYMS protein, we next examined whether SIN1 depletion affects cellular sensitivity to short and long-term treatment with pyrimidine analogs. Compared to Sh control cells, SIN1 depletion sensitizes the cells to FuDR-mediated apoptosis, as seen with the increase in the percentage of SubG1 cells. Together, these results demonstrate that SIN1 is required for cellular response to stress and protects cells from 5-Fu and FuDR.

## Discussion

Resistance to 5-Fluorouracil (5-FU) is directly linked to elevated levels of its target protein, thymidylate synthase (TYMS). Several mechanisms controlling TYMS levels have been proposed. It was first suggested that FdUMP interaction with TYMS releases its RNA, which in turn promotes translation [10-12]. Using colorectal cells, Varghese et al. reported that forkhead box transcription factor, FOXM1, could be involved in positive regulation of TYMS [29]. Despite being controversial, data from Intuyod et al. showed that FOXM1 knockdown had only a moderate effect on TYMS levels [30]. This raises the question of which factor directly controls TYMS levels and dynamics. In the current study, we provide the first mechanistic evidence that TYMS protein dynamic is mainly regulated via a mechanism that involves translation rather than transcription in response to 5-fluorouracil (5-FU) and its derivative, fluorodeoxyuridine (FuDR). Our findings challenge the traditional view that TYMS regulation in the context of 5-FU treatment is mainly regulated at the transcriptional level. Instead, we show here that in breast cancer cells, 5-FU and FuDR treatment results in a biphasic response in TYMS protein levels degradation within the first two hours, followed by a significant accumulation at 24 hours. Importantly, these changes occur independently of transcriptional changes, as TYMS mRNA levels remain stable during the initial phase and increase modestly at later time points.

The absence of changes in TYMS mRNA in the early phase of treatment suggests that the observed decrease in TYMS protein levels are not a result of transcriptional regulation, but rather changes in translation. This hypothesis was further supported by metabolic labelling experiments, which demonstrated a significant decrease in TYMS translation. It is very well established that mTORC1 plays a crucial role in regulation of translation. However, emerging evidence also suggests a role for mTORC2 in this process. Indeed, mTOR2 association with the ribosome is required for cotranslational phosphorylation of AKT, its stability, and signalling [31-32]. Our data show that 5-FU treatment negatively alters mTORC1 activity as seen by decreased P70S6 kinase phosphorylation at both early and later time points, suggesting that mTORC1 is not involved in TYMS dynamics. Using thermal protein profiling, we next identified mTORC2 component SIN1 among the top proteins that showed changes in thermal stability after two hours of treatment with 5-FU. Although the mechanism behind this alteration is unknown, our data provide direct evidence that this protein controls TYMS protein levels and dynamics. Indeed, SIN1 knockdown significantly alters TYMS RNA and protein levels under both basal conditions and upon 5-FU or FUDR treatment. These findings point to translation as the dominant regulatory mechanism controlling TYMS protein in response to these chemotherapeutic agents.

Although a comprehensive analysis of global and selective translational changes following 5-FU treatment is lacking, our results are consistent with recent findings implicating translational changes in response to 5-FU as a key component of the cellular response to 5-FU [33]. In our study using breast cancer cells, we observed a decrease and an increase in mTORC1 and mTORC2, respectively. In contrast, a previous study in colorectal cancer cells reported increased mTORC1 activity upon 5-FU treatment. This discrepancy could be attributed to differences in stress response pathways and translational control mechanisms between different cancer types. Nevertheless, both studies consistently highlight the impact of 5-FU on factors that affect translation machinery linked to pro-survival pathways. Together, these observations suggest that modulation of translation is a critical determinant of cell fate in response to 5-FU across diverse cancer types. Therefore, co-targeting translation machinery with 5-FU could be a strategy to enhance therapeutic efficacy, potentially overcoming resistance mechanisms.

## Materials and Methods

### Reagents

DMEM High Glucose (4.5 g/l), with L-Glutamine, Sodium Pyruvate (Capricorn Scientific, #DMEM-HPA), Fetal Bovine Serum (Sigma Aldrich, # F7524-500ML), Penicillin-Streptomycin (Thermo Fisher Scientific, #15140122). The following inhibitors were used: 5-fluorouracil (Sigma Aldrich, #F6627), 5-Fluoro-2’-deoxyuridine (Thermo Scientific Chemicals, #L16497.ME), 5-fluorouridine (Thermo Scientific Chemicals, #J62083.03), Rapamycin (Thermo Scientific Chemicals, #J62473.MC).

### Cell culture

HEK293T, MDA-MB-231, and MCF7 cells were generously provided as a gift from Dr. Anna Marusiak, IMol Institute. The cells were cultured in DMEM supplemented with 10% fetal bovine serum (FBS), 100 U/mL penicillin, and 100 U/mL streptomycin. Cells were maintained at 37 °C in a humidified, 5% CO2 incubator. The concentrations and duration of treatment of cells with the above inhibitors were all described in the figure legends. In each experiment, control cells were treated with an equivalent volume of solvent (DMSO).

### Proteome Integral Solubility Alteration Analysis of 5-Fu

PISA assay was carried out as described previously [23]. Briefly, MDA-MB-231 cells were seeded into a 10 cm culture dish at a confluence of 80 %. Twenty-four hours later, cells were treated for two hours with 200 μM 5-FU or DMSO as a control, three biological replicates for each condition. After treatment, cells were detached using Trypsin-EDTA solution (Thermo Fisher Scientific, *#*15090046) and washed twice with phosphate-buffered saline (PBS). Cell pellets were collected and resuspended in 500µL of PBS. After homogenization, each biological replicate was split into eight 0.2 mL PCR tubes, 60μl/tube, one for each temperature point. The following temperatures were used for PISA (43°C, 44.3°C, 46.2°C, 48.5°C, 51.8°C, 54.3°C, 55.9°C, 57°C). Thermal treatment was carried out for 3 minutes using a C1000 Touch Thermal Cycler (BioRad). Samples were then incubated at room temperature for 5 min. Cell lysates were obtained by freeze-thaw cycles in liquid nitrogen three times and then thawing at 30 °C. Aliquots of the same biological replicates were recombined. Soluble cell lysates were obtained after two rounds of centrifugation at 20000g for 20 min at 4 °C. Soluble fractions from each condition were collected, Protein concentrations were determined using the Pierce™ BCA Protein Assay Kit (Thermo Scientific™ #23225), and BSA standards were used in parallel according to the manufacturer’s instructions. Twenty μg of each sample was digested using trypsin overnight at 37 °C with vortexing at 1000 rpm. Digested samples were then labeled with TMT as follows. First, three layers of the resin C18 mesh were packed inside 200 µL pipette tips, which were inserted onto the lid of the Eppendorf tube called STAGE Tips-Column. Columns were first conditioned using 150 µL methanol at 1200 g for 2 minutes, then washed with 100 µL (50% acetonitrile/0.1% formic acid) at 1200 g for 2 minutes. The resin was then equilibrated twice with 150 µL 0.1% formic acid at 1200 g for 2 minutes.

Ten 10 µg equivalent volumes of the digested samples were loaded onto the STAGE Tips-Column and centrifuged at 1200 g for 2 minutes. STAGE Tips-Columns were washed twice with 150 µL of 0.1% formic acid (1200 g, 2 min).

For peptide labeling, TMTs were first resuspended in 2 µL acetonitrile, followed by the addition of 200 µL of freshly prepared 50 mM HEPES, pH 8. Peptide labeling was done by loading 200 µL TMT solution onto STAGE Tips, followed by 300 g centrifugation for 10 minutes. Labeled peptides were washed 3 times with 150 µL of 0.1% formic acid (1200 g, 2 min). Labeled peptides were eluted from the resin using 60 µL 60 % acetonitrile (1200 g, 2 min). A volume of 55 µL was taken from each sample and combined, then dried in a centrifugal vacuum concentrator at 40 °C (SpeedVac). Dried samples were reconstituted in 0.1% Trifluoroacetic acid and fractionated using Pierce High pH Reversed-Phase Peptide fractionation kit (# 84868, Thermo Scientific™), following manufacturer instructions. Fractionated peptides were dried in a centrifugal vacuum concentrator at 40 °C (SpeedVac), reconstituted in 0.1% formic acid, and measured using LC-MS/MS.

### LC-MS/MS measurements and data analysis

The peptide fractions were resuspended in 0.1% TFA and 2% acetonitrile in water. Chromatographic separation was performed on an Easy-Spray Acclaim PepMap column (50 cm length × 75 µm inner diameter; Thermo Fisher Scientific) at 55°C by applying 120 min acetonitrile gradients in 0.1% aqueous formic acid at a flow rate of 300 nl/min. An UltiMate 3000 nano-LC system was coupled to a Q Exactive HF-X mass spectrometer via an easy-spray source (all Thermo Fisher Scientific). The Q Exactive HF-X was operated in TMT mode with survey scans acquired at a resolution of 60,000 at m/z 200. Up to 15 of the most abundant isotope patterns with charges 2-5 from the survey scan were selected with an isolation window of 0.7 m/z and fragmented by higher-energy collision dissociation with normalized collision energies of 32, while the dynamic exclusion was set to 35 s. The maximum ion injection times for the survey scan and dual MS (MS/MS) scans (acquired with a resolution of 45,000 at m/z 200) were 50 and 120 ms, respectively. The ion target value for MS was set to 3e6, and for MS/MS, it was set to 1e5, and the minimum AGC target was set to 1e3.

The data were processed with MaxQuant v. 1.6.17.0, and the peptides were identified from the MS/MS spectra searched against Uniprot Human Reference Proteome (UP000005640) using the built-in Andromeda search engine [34-36]. Reporter ion MS2-based quantification was applied with reporter mass tolerance = 0.003 Da and min. reporter PIF = 0.75. Cysteine carbamidomethylation was set as a fixed modification and methionine oxidation, glutamine/asparagine deamination, as well as protein N-terminal acetylation were set as variable modifications. For in silico digests of the reference proteome, cleavages of arginine or lysine followed by any amino acid were allowed (trypsin/P), and up to two missed cleavages were allowed. The FDR was set to 0.01 for peptides, proteins, and sites. Match between runs was enabled. Other parameters were used as pre-set in the software. Reporter intensity corrected values for protein groups were loaded into Perseus v. 1.6.10 [37]. Standard filtering steps were applied to clean up the dataset: reverse (matched to decoy database), only identified by site, and potential contaminant (from a list of commonly occurring contaminants included in MaxQuant) protein groups were removed. Reporter intensity corrected values were log2 transformed, and protein groups with values across all samples were kept. The values were normalized by median subtraction within TMT channels. To determine proteins whose thermal stability was affected by the supplementation of 5-FU compared to the vehicle control, Student’s t-test (2-sided, permutation-based FDR = 0.05, S0 = 0.1, n = 3) was performed. The table was exported from Perseus and formatted to its final form in Microsoft Excel 2016.

### Western Blotting

Vehicle and control-treated cells were first washed with PBS. Protein lysates were prepared using lysis buffer containing 50 mM Tris-HCl, pH 8.0, 150 mM NaCl, 1.0% NP-40, 0.1% SDS, 25 units/ml Benzonase and 100 µM PMSF (#36978, Thermo Scientific™). Total protein extracts were isolated by centrifugation at 12000g for 10 minutes at 4°C. Protein concentrations were determined using a BCA assay (Thermo Scientific™ #23225). Equal amounts of proteins (10–20 µg) were then resolved using SDS-PAGE and transferred to a Polyvinylidene fluoride (PVDF) membrane. Upon transfer, membranes were blocked with tris-buffered saline containing 0.05 % Tween 20 supplemented with 5 % non-fat milk for 1 hour, followed by overnight incubation in primary antibodies. Subsequently, membranes were washed three times with TBST, followed by incubation in an appropriate dilution of secondary antibody at room temperature for 1 hour. Antibody dilutions were prepared following the manufacturer’s instructions. The chemiluminescence signal was detected using Amersham ImageQuant 800 systems. The following antibodies were purchased from invitrogen: phospho-AKT1 (Ser473) # 700256; AKT Pan # MA5–14916, Phospho-p70 S6 Kinase (Thr389) #710095, P70 S6 Kinase #MA5-15141, P21 Monoclonal Antibody (R.229.6) # MA5-14949, GAPDH # MA5-15738. SIN1 antibody was purchased from Bethyl Laboratories #A300-910A-T.

### Packaging shRNA-Encoding Lentivirus

HEK293T cells were plated overnight in 6 well plates at 50% confluency. The medium was replaced before transfection. A mixture containing Packaging plasmid psPAX2, 1 μg (addgene#12260), Envelope plasmid VSV-G, 0.2 μg (addgene #12259), and 1μg of each sh-RNA (Control and targeting Sh-RNA) were resuspended in 100 μL of Opti-MEM (Gibco, #31985062). Second mixture containing 6.6 µg polyethylenimine (PEI) (Sigma Aldrich, #913375) and 100 μL of Opti-MEM was prepared in a separate tube. The two mixtures were combined, vortexed, and incubated for 20 minutes at RT. The mixture was then added to each well dropwise. The medium was replaced after 4 hours. Forty-eight hours later, lentiviral supernatants were collected and centrifuged for 5 minutes at 12000g, then stored at −80°C until further use. The following shRNAs were used: SIN1 (shRNA1: Addgene #13483, and shRNA2: Addgene #13484). ShRNA control Empty (Sigma Aldrich, # SHC001).

### Lentivirus infection and selection

MCF7 and MDA-MB-231 cells were plated overnight in 6 well plates at 50% confluency. For virus infection, 1 mL of fresh medium was mixed with 1 mL of lentiviral supernatant, then directly added to the cells and left overnight. Upon medium change, cells were incubated for an additional 24 hours. The selection was carried out using puromycin (Sigma Aldrich, # P4512). The following concentrations were used: MCF7 3µg/ml, MDA-MB-231 2 µg/ml) for 96 hours. Real-time PCR and western blot were used to verify the Knockdown efficiency.

### RNA Isolation

Vehicle and control-treated cells were washed with PBS and resuspended in appropriate volumes of TRI Reagent (Thermo Fisher Scientific, #AM9738). RNA isolation was carried out according to the manufacturer’s instructions. Briefly, cells were lysed in TRI Reagent by incubating for 5 min at room temperature. The mixture was then supplemented with an appropriate volume of chloroform, followed by vigorous shaking for 20 seconds. Samples were incubated at room temperature for an additional 5 minutes and then centrifuged at 15,000 × g for 15 min at 4°C. The aqueous containing RNAs were transferred to a new tube. RNA precipitation was carried by adding one volume of isopropanol and incubating at −20°C for 1 hour, followed by centrifugation at 12,000 × g for 20 min at 4°C. The RNA pellet was washed with 75% ethanol and centrifuged at 12,000 × g for 10 min at 4°C. RNA concentrations were determined, and cDNA synthesis was carried out as described below.

### cDNA synthesis and quantitative real-time PCR

RNA samples prepared above were used for cDNA synthesis using the High-Capacity cDNA Reverse Transcription Kit (#4368814, ThermoFisher), 500ng of RNA per reaction. Then, Real-time PCR was carried out using PowerUp™ SYBR™ Green Master Mix (ThermoFisher, #A25776) according to the manufacturer’s instructions. Briefly, a mixture containing 10 ng of cDNA, 7 μL SYBR™ Green Master Mix, 300 nM forward and reverse primers, and appropriate volumes of nuclease-free water was prepared and analyzed in duplicates using a Light Cycler 480 Real-Time PCR System. The following steps were used: First, an activation step at 95°C for 2 min, followed by 40 cycles at 95°C for 30 s, 61°C for 30 s, and 72°C for 30 s. A melting curve analysis of the PCR products was performed to verify the specificity of each pair of primers. Expression levels of the indicated genes were analyzed and quantified relative to Gapdh. The following primers were used:

Forward-TYMS-CTGCTGACAACCAAACGTGTG,
Reverse-TYMS-GCATCCCAGATTTTCACTCCCTT,
Forward-SIN1-GGTGGACACCGATTTCCCC,
Reverse-SIN1-CGCTTCACTGCCTTCAGTAAGA.
Forward-GAPDH-GGAGCGAGATCCCTCCAAAAT,
Reverse-GAPDH-GGCTGTTGTCATACTTCTCATGG,

### Drug Affinity Responsive Target Stability (DARTS)

DART assay was carried out as described previously [15]. Briefly, cell lysate was prepared using NP-40 lysis buffer containing 50 mM Tris-HCl pH 8.0, 150 mM NaCl, 1% NP-40, and 1 mM PMSF. Lysates were then cleared by centrifugation at 10000 g for 10 minutes at 4°C. Protein concentrations were first determined using a BCA assay (Thermo Scientific™ #23225). Equal amounts of lysates (100–150 µg) were treated with DMSO or the indicated inhibitors and incubated on a rotating shaker for 30 minutes at RT. Mixtures were collected by short pulse centrifugation, supplemented with control solution or pronase (Roch, #10165921001), and digested for 15 minutes at RT. Digestion was stopped by adding Laemmli Sample Buffer and incubating at 95 °C for 5 min. Samples were then analyzed using a regular western blot and stained with the indicated antibodies as described above.

#### Metabolic labeling of proteins in MDA-MB-231 cells with β-ethynylserine (βES)

Metabolic labeling was carried out as described previously [38-39]. An equal number of MDA-MB-231 cells were seeded in a T-75 flask and grown overnight in complete medium. Labeling experiments were performed by adding 1 mM **βES** at 37 °C in a complete medium for 60 min. Control samples were simultaneously pre-treated with 10 µg/ml cycloheximide to block translation. One hour later, cells were treated with DMSO, 50 µM FuDR, or 50 µM FuR for an additional 2 hours. At the end of incubation, cells were washed with PBS and resuspended in 200 μL of lysis buffer containing 2 % SDS in PBS. Samples were sonicated using a probe sonicator (1× 10 seconds) at room temperature

#### Bioorthogonal conjugation of βES-labeled proteins to Azide-PEG3-biotin in cell lysate, enrichment of biotinylated proteins, and targeted detection by Western blotting

Lysates prepared after labeling cells with β-ethynylserine above were used as follows. Lysates were diluted with 200 μL PBS, then supplemented with 40 mM iodoacetamide and incubated at room temperature for 40 min under shaking with a thermomixer (900-1000 rpm at room temperature). Proteins were then precipitated with 400 μL methanol and 100 μL chloroform, then collected by centrifugation (4000× g, 3 min) at RT. The protein pellet was washed twice with methanol and dissolved in 50 μL PBS containing 2 % SDS, followed by 150 μL PBS (0.5 % SDS final).

Copper-catalyzed azide-alkyne click (CuAAC) reaction was performed for 1 hour at RT by adding 14 µL CuAAC reaction mixture to the 200 μL proteins. The final concentrations of CuAAC in the reaction mixture are as follows: 1 mM CuSO4, 200 μM THPTA, 1 mM TCEP, and 100 μM Azide-PEG3-biotin. The click reaction was stopped by adding 2 μL of 500 mM EDTA. Proteins were then precipitated with 1 volume of methanol and ¼ volume of chloroform, then collected by centrifugation (4000× g, 3 min) at RT. The protein pellet was washed twice with methanol and dissolved in 100 μL PBS containing 2 % SDS, and diluted with 300 μL of PBS. Finally, enrichment of biotine-βES-protein conjugates was achieved using Pierce™ NeutrAvidin™ Agarose beads (Thermo Scientific™, #29201), following the manufacturer’s instructions. Bound proteins were eluted using SDS sample loading buffer, separated using SDS-PAGE, transferred onto nitrocellulose membrane, and probed with protein-specific antibodies TYMS, AKT, and PARP1.

#### Cell cycle analysis (SubG1 measurement)

MDA-MB-231 cells were harvested following 72 h treatment with DMSO or 50 µM FuDR. Cells were first trypsinized, washed twice with PBS, and fixed in 70% ethanol at −20°C for 24 hours. Fixed cells were then resuspended in 400 mL of PBS supplemented with 0.2 mg/ml DNase-free RNase A (Thermo Scientific™, EN0531), 10 µg/ml propidium iodide (Invitrogen™, P1304MP), and incubated at 4°C for 2 hours protected from light. Stained cells were analyzed using a name flow cytometer and software for DNA content determination.

### Statistical analysis

Statistical analysis was performed using GraphPad Prism (http://www.graphpad.com). A minimum of three biological replicates were carried out for each experiment. An unpaired, two-sided test was used to compare the means of two groups. One-way ANOVA with Dunnett test post hoc test was applied to compare the means of three or more groups. All results in the graphs are expressed as means ± SEM. *P* < 0.05 was considered to be significant.

---

**Supplementary Figure 1.**

**(A-B)** Lysates from MDA-MB-231 and MCF-7 cells treated with the indicated concentrations of 5-FU for 2 hours. **(C-D)** Bar graphs showing TYMS protein levels under conditions obtained in A and B, respectively. The data are presented as the means ±SEMs. ANOVA, multiple comparisons: Dunnett test. MDA-MB-231: (F (3, 8) = 11.97, P˂0.005. MCF7: F (3, 8) = 24.26, P˂0.001).

**Supplementary Figure 2.**

**(A)** Lysate from MDA-MB-231 cells treated with 50µM FuR for the indicated durations and then stained with the indicated antibodies. **(B)** Bar graphs showing TYMS protein levels under conditions obtained in A. The data are presented as the means ±SEMs. ANOVA, multiple comparisons: Dunnett test. F (3, 8) = 11.94, P˂0.005. (**C**) Lysate from MDA-MB-231 cells treated with 50µM FuR for 24 hours and stained with the indicated antibodies. 3 separate biological replicates. (**D**) Bar graph showing TYMS protein levels under different conditions obtained in C. The data are presented as the means ±SEMs. Unpaired T test, **, P˂0.01. **(E)** Results of DART assay. MDA-MB-231 lysate treated or not with 20µM FuDR or 20µM FuR for 30 min, then digested for 10 min with pronase, and then stained for the indicated antibodies. (**F**) Bar graph showing TYMS protein levels under different conditions obtained in E. The data are presented as the means ±SEMs. ANOVA, multiple comparisons: Dunnett test. F (3, 8) = 49.34, P<0.0001.

**Supplementary Figure 3.**

(**A)** Lysates from MCF-7 cells pre-treated with 2 µM Mg132 or 100 µM chloroquine for 2 hours, followed by 100µM 5-FU for 2 hours. **(B)** Bar graphs showing TYMS protein levels under different conditions obtained in A. The data are presented as the means ±SEMs. ANOVA, multiple comparisons: Dunnett test. F (3, 8) = 60.63, P<0.0001.

**Supplementary Figure 4. 5-Fu and its derivative alter mTORC1/2 activity.**

**(A-B, E-F)** Lysates from MCF-7 and MDA-MB-231 cells treated respectively with 100 µM 5-Fu or 50 µM FuDR for the indicated durations and then stained with the indicated antibodies. **(C-D, G-H)** Bar graphs showing the results of densitometric analysis of AKT1 and P70SK phosphorylation under different conditions obtained in A-B and E-F, respectively. The data are presented as the means ±SEMs. ANOVA, multiple comparisons: Dunnett test. **5-Fu:** MDA-MB-231: p-AKT1 (F (3, 8) = 54.98, P<0.0001), p-P70SK (F (3, 8) = 10.19 P<0.005). MCF7: p-AKT1 (F (3, 8) = 39.42, P<0.0001), p-P70SK (F (3, 8) = 9.914, P<0.005). **FuDR**: MCF7: p-AKT1 (F (3, 8) = 3.547, P=0.0676), p-P70SK (F (3, 8) = 68.21, P<0.0001). MDA-MB-231: p-AKT1 (F (3, 8) = 3.204, P=0.0835), p-P70SK (FF (3, 8) = 4.933, P<0.05).

**Supplementary Figure 5.**

**(A)** Lysates from MDA-MB-231 cells treated or not with 50nM (GDC-0980), 50nM wortmannin, 50nM Rapamycin, 50nM autophagy inhibitor or 5nM Bafilomycin A during 24 hours stained with the indicated antibodies. Two separate biological replicates were shown.

**Supplementary Figure 6.**

**(A-C)** RNA and protein levels of SIN1 in MDA-MB-231 (A-B) and in MCF7 (C) cells stably transfected with the shRNA control (Ctrl) or shRNA against SIN1. Bar graph showing the qPCR results. Data are presented as the means ± SEMs. Unpaired t-test, *, P˂0.01. (C–D)

