## supplementray figures for "Mechanistic Insights into TYSM Protein Regulation by mTORC2 in Response to Chemotherapy"

### Supplementary Figure 1

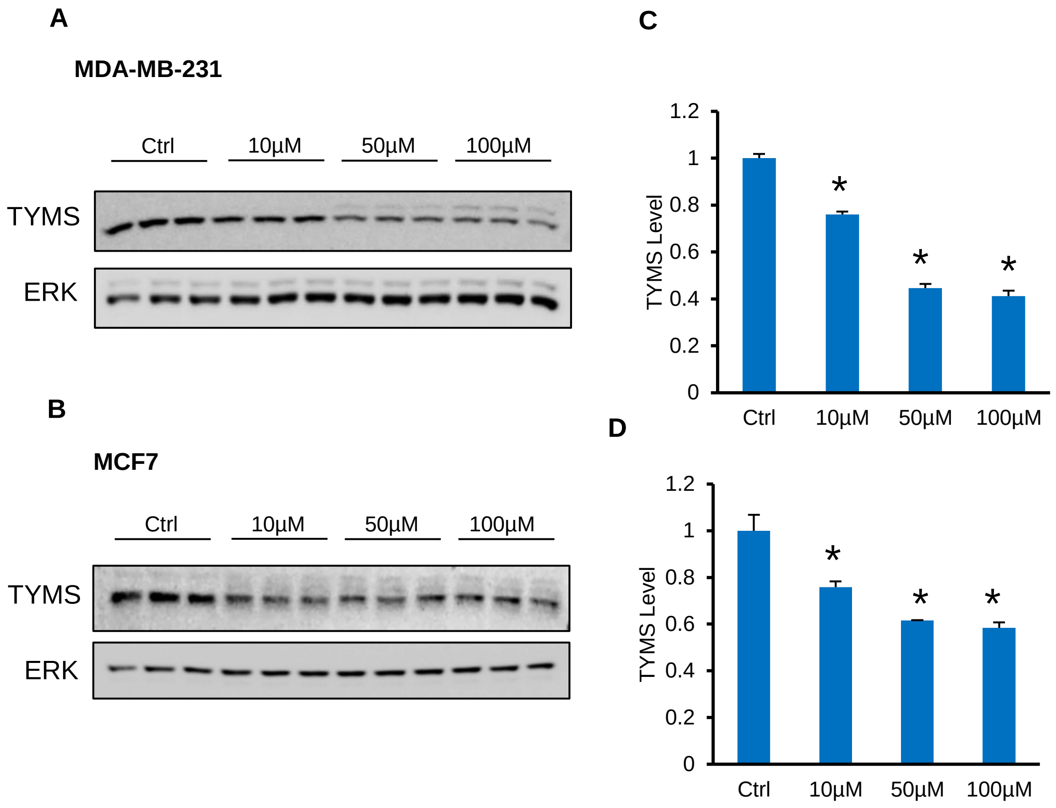

### Supplementary Figure 2

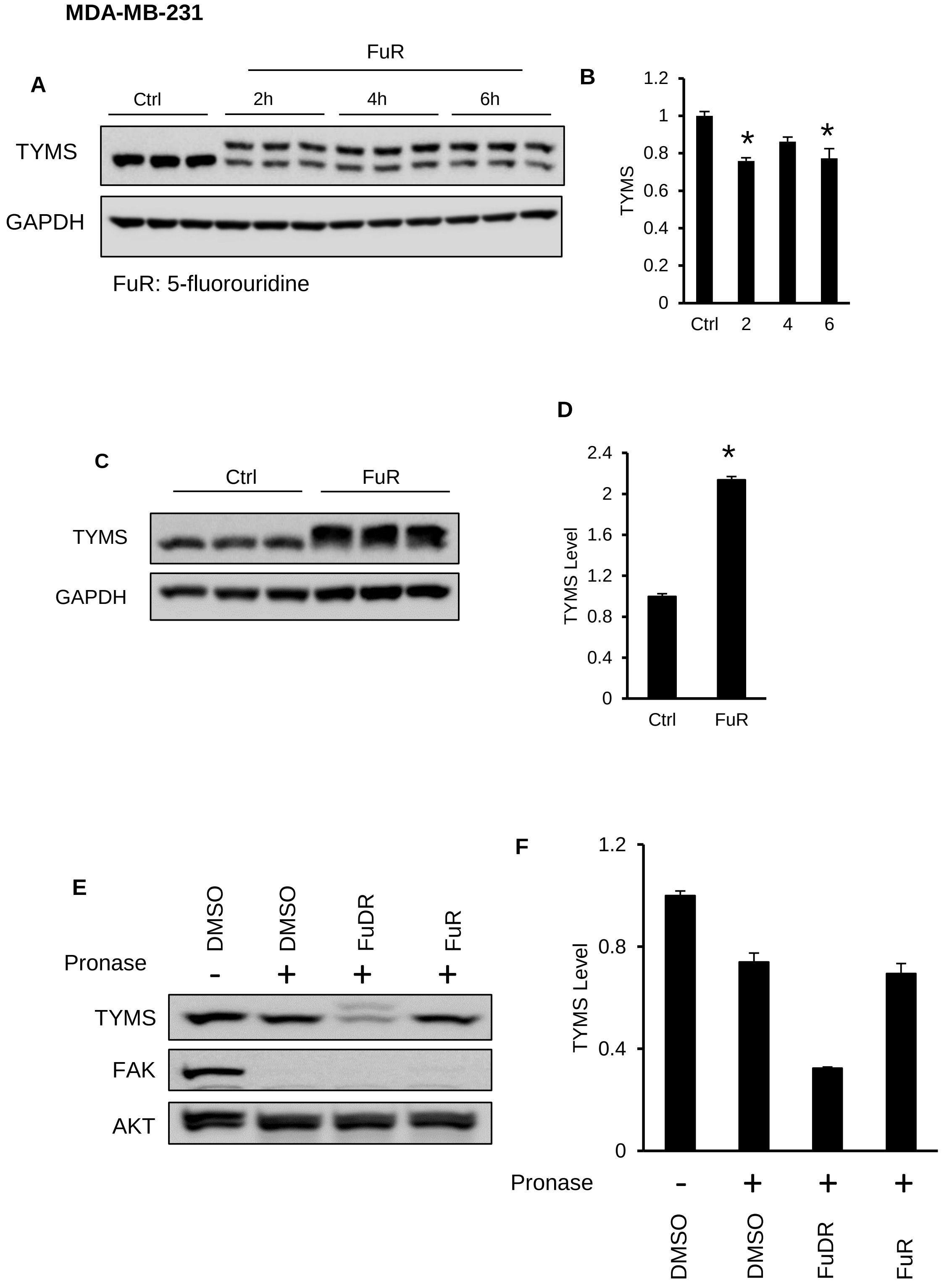

### Supplementary Figure 3

A

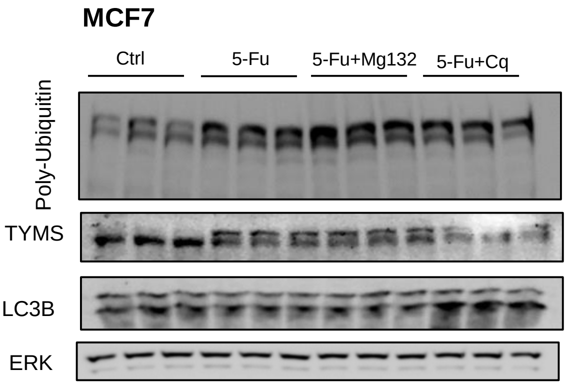


B

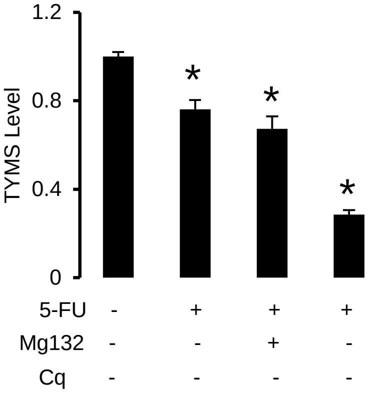

### Supplementary Figure 4

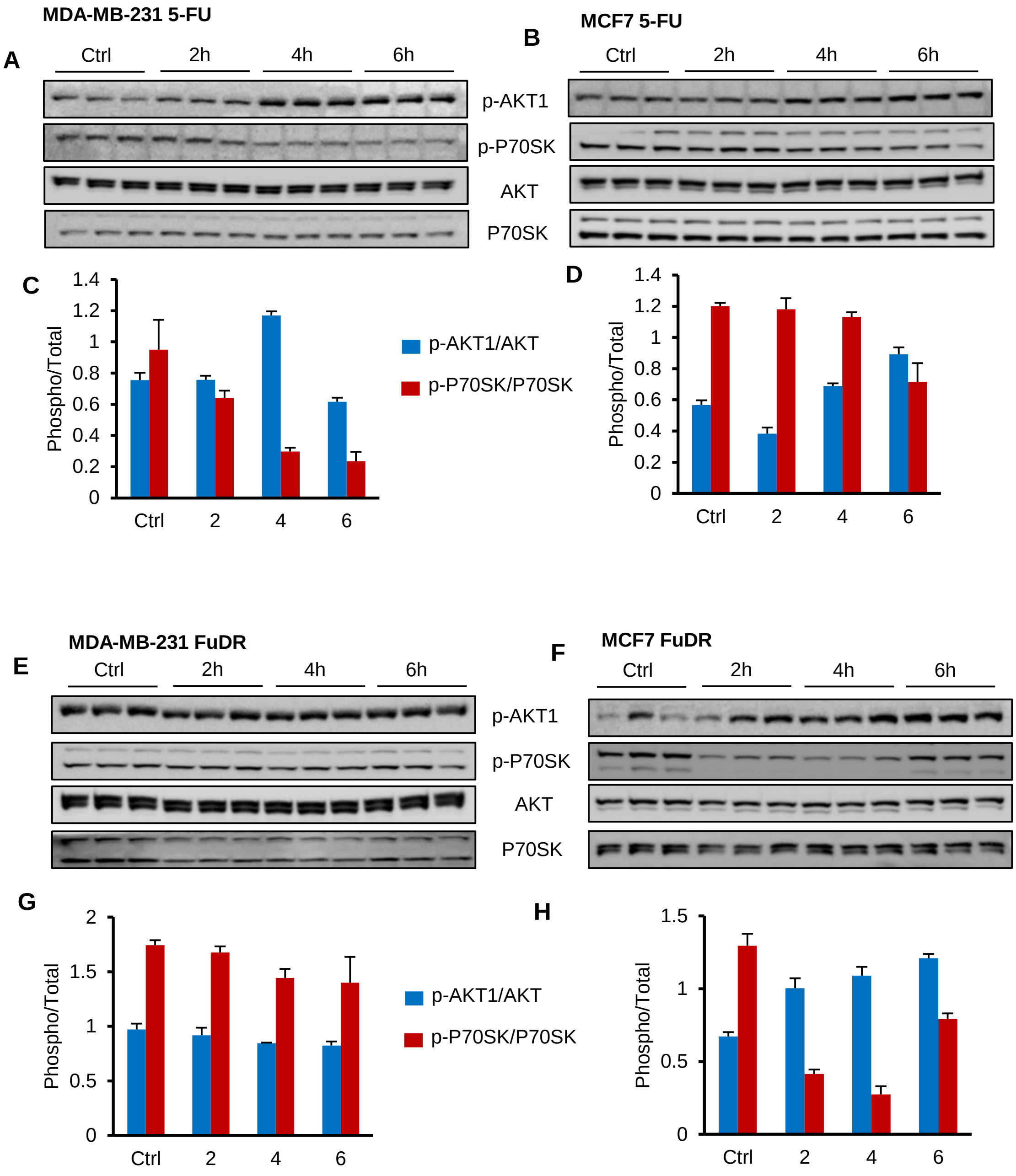

### Supplementary Figure 5

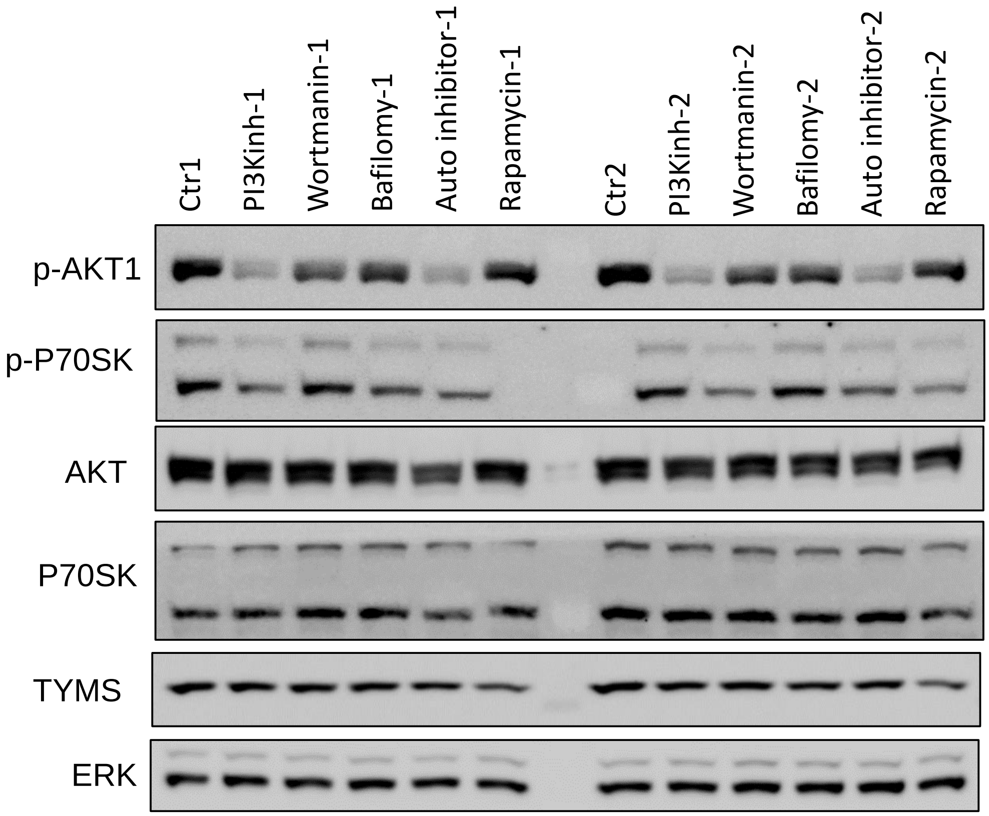

### Supp Figure 6

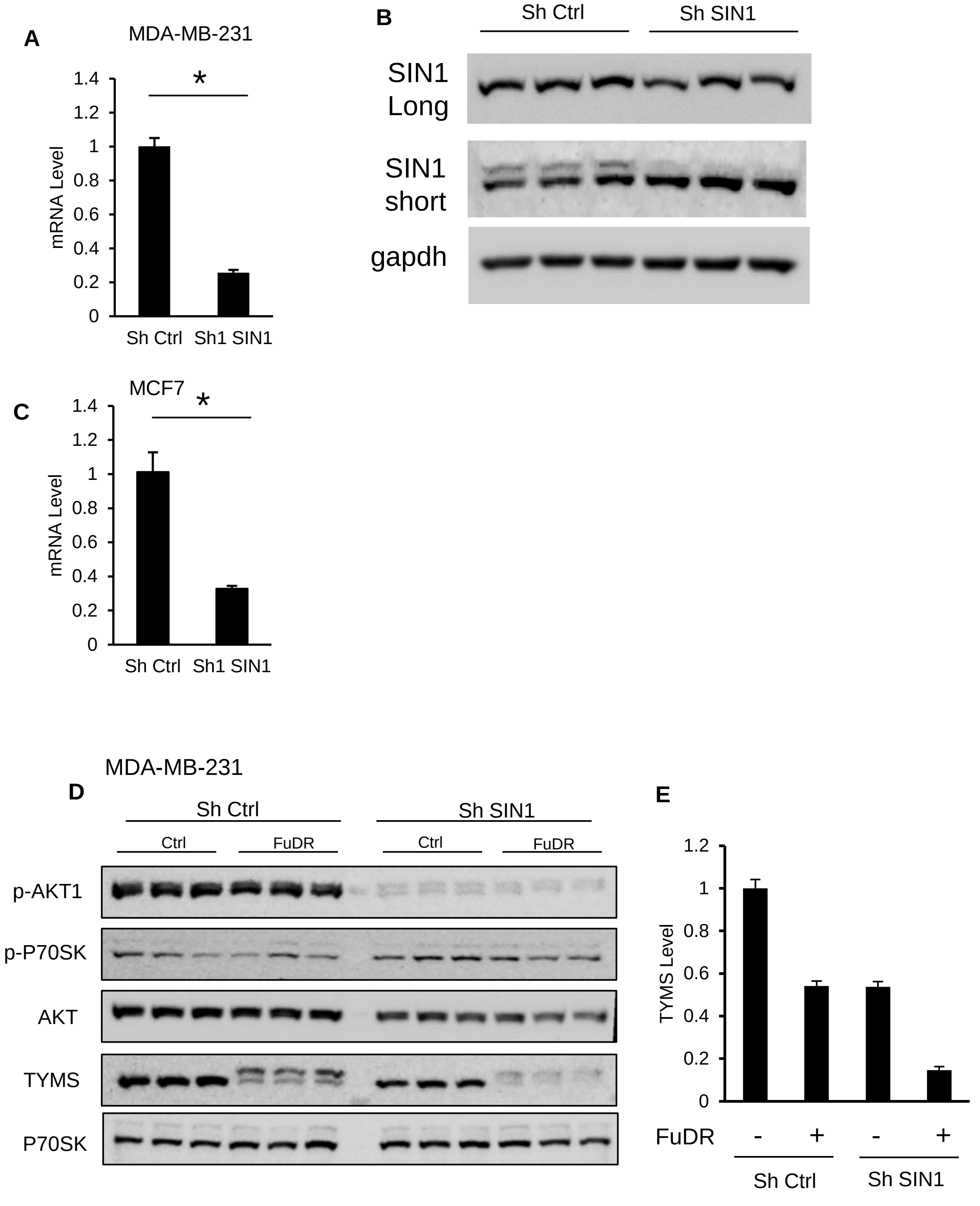
